# Mixture of Experts Enable Efficient and Effective Protein Understanding and Design

**DOI:** 10.1101/2024.11.29.625425

**Authors:** Ning Sun, Shuxian Zou, Tianhua Tao, Sazan Mahbub, Dian Li, Yonghao Zhuang, Hongyi Wang, Xingyi Cheng, Le Song, Eric P. Xing

**Affiliations:** GenBio AI; Mohamed bin Zayed University of Artificial Intelligence; Carnegie Mellon University; University of Washington

## Abstract

Proteins play a fundamental role in life. Understanding the language of proteins offers significant potential for gaining mechanistic insights into biological systems and introduces new avenues for treating diseases, enhancing agriculture, and safeguarding the environment. While large protein language models (PLMs) like ESM2-15B and xTrimoPGLM-100B have achieved remarkable performance in diverse protein understanding and design tasks, these models, being dense transformer models, pose challenges due to their computational inefficiency during training and deployment. In this work, we introduce AIDO.Protein, a pretrained module for protein representation in an AI-driven Digital Organism [1]. AIDO.Protein is also the first mixture-of-experts (MoE) model in the protein domain, with model size scales to 16 billion parameters. Leveraging a sparse MoE architecture with 8 experts within each transformer block and selectively activating 2 experts for each input token, our model is significantly more efficient in training and inference. Through pre-training on 1.2 trillion amino acids collected from UniRef90 and ColabfoldDB, our model achieves state-of-the-art results across most tasks in the xTrimoPGLM benchmark. Furthermore, on over 280 ProteinGym Deep Mutational Scanning (DMS) assays, our model achieves nearly 99% of the overall performance of the best MSA-based model and significantly outperforms the previously reported state-of-the-art models that do not utilize MSA. We also adapted this model for structure-conditioned protein sequence generation tasks and achieved new SOTA in this domain. These results indicate that AIDO.Protein can serve as a strong foundation model for protein understanding and design. Models and codes are available through ModelGenerator in https://github.com/genbio-ai/AIDO and on Hugging Face.

## 1 Introduction

As the end product of genes, proteins serve as the workhorses of life, carrying out most of the biological functions within the cell. They act as biological catalysts, provide structural support to cells and tissues, facilitate the transport of molecules across cell membranes and within cells, recognize and neutralize foreign substances like pathogens, transmit signals that regulate cellular processes, etc. To understand the function of a protein, a line of work follows the sequence-structure-function relationship, studying the structure first in order to understand its function since 3D structure is the active form of a protein [2]. Since protein structures are quite scarce, another line of work goes directly from sequence to function, aiming to determine the function given sequence-only information [3, 4]. Under both directions, we can see a common need to understand the language of protein sequences. Understanding the language of proteins is crucial to advancing genetic research and accelerating drug discovery. For example, it can help design enzymes that metabolize plastic waste or hydrolyse polluting toxins. It can also help create vaccines in a timely fashion during a pandemic.

Recent advances in artificial intelligence, especially the large language modeling technologies, offer promising avenues toward this goal. The huge success of large language models (LLMs) in natural language processing (NLP) inspire researchers to apply self-supervised pre-training in the protein domain, using protein sequences without any structural and functional labels [5, 6, 7, 8, 9, 10, 11, 12, 13, 3, 14, 15, 4]. These protein language models have demonstrated remarkable performance in a diverse array of tasks, including protein structure prediction, protein function prediction, and protein sequence design. For example, ESMFold based on protein language models [3] achieves atomic accuracy in protein structure prediction, reaching near AlphaFold2 [2] performance. xTrimoPGLM [4], a PLM at the scale of 100B parameters, obtains superior performance in diverse taks of protein function prediction. One of the key drive of performance is the scale of the PLM and we can see a clear trend that larger models have better performance. However, the computation efficiency will decrease when PLMs grow larger since running larger models is computationally more expensive. Exisiting work focus on pushing the performance of PLMs by scaling up model size without concerning much on the computation efficiency. Here we ask: *can we maintain/push the performance of a PLM while keeping it efficient during training and inference?*

We resort to sparse expert models for a potential solution. Sparse expert models are neural networks in which a subset of parameters is divided into “experts”, each having a distinct weight. The models route input examples to specific expert(s) weights during training and inference. As a result, each example only interacts with a subset of the network parameters, different from dense models. Because only a fraction of the experts are used for each example, the amount of computation may remain small relative to the total model size [16]. Significant work has been done to investigate sparse expert models, of which, Mixture-of-Experts (MoE) stands out. Integrated into Transformer, MoE has become a strong counterpart of dense transformer models in the NLP domain [17, 18, 19].

In this work, we explore pre-training the first MoE model in the protein domain, different from all existing PLMs which adopt dense transformer architecture. We present AIDO.Protein, a MoE model at the scale of 16 billion parameters, pre-trained on 1.2 trillion tokens collected from UniRef90 and ColabfoldDB. During training and inference, each input token is processed by 4.5 billion parameters, using only 28% of the total number of parameters. We evaluate our model in a wide range of tasks, including 18 diverse tasks from xTrimoPGLM benchmark and 283 protein fitness prediction tasks from ProteinGym DMS benchmark. Experiment results show that AIDO.Protein achieves strong performance across the board while being more computational efficient. We further leverage AIDO.Protein for protein inverse folding and find that it outperforms previous SOTA methods, such as ProteinMPNN [20] and LM-Design [21]. These results demonstrate the effectiveness of AIDO.Protein in protein sequence understanding and design, providing the community with a new powerful and efficient protein foundation model.

## 2 Related work

### 2.1 Protein language model

Inspired by the huge success of large language models (LLMs) in the natural language processing (NLP) domain, in recent years, there has been a line of work applying LLM technologies in the protein domain [5, 6, 7, 8, 9, 10, 11, 12, 13, 3, 14, 15, 4].By pre-training on protein sequence databases, these protein language models have gain remarkable abilities in extracting biological meaningful representations for various downstream tasks, including protein structure and function predictions. In particular, ESM2 series, which scales up to 15 billion parameters, achieves atomic accuracy in protein structure prediction, demonstrating the effectiveness of large protein language model [3]. Recently, Chen et al. (2024) [4] pre-train a protein language model named xTrimoPGLM that conatains 100 billion parameters. It achieves superior performance on diverse tasks over ESM2-15B model, further showcasing the effectiveness of scaling in the protein domain. However, these large protein language models are all dense transformer models, making finetuning for downstream tasks computational intensive especially for ESM2-15B and xTrimoPGLM-100B. In addition, the current largest opensource protein language model before our model is ESM2-15B since xTrimoPGLM-100B is not publicly available.

### 2.2 Mixture of experts

The scale of a model is one of the most important factors for better model quality. Given a fixed computing budget, training a larger model for fewer steps is better than training a smaller model for more steps [22]. Mixture of Experts (MoE) enable models to be pre-trained with far less compute, which means the model or dataset size can be dramatically scale up with the same compute budget as a dense model. It is a sparse neural network which leverages multiple expert networks, with a gating mechanism to select the most relevant experts for each input [23, 24]. This approach has gained prominence in large language models, where MoE layers replace dense MLP layers in transformers, allowing models to scale more efficiently by activating only a subset of experts for each token [16]. Key advancements include the GShard [17] and Switch Transformer [18], which improved training stability and efficiency through selective expert activation and load balancing strategies. The MoE design has also been successfully applied in vision transformers, with models like V-MoE [25] achieving comparable performance to dense models with significantly reduced computational costs. These developments highlight MoE’s potential in both natural language processing and computer vision, offering scalable solutions for complex tasks. Recently, a powerful MoE model called Mixtral 8×7B [19] outperforms Llama 2 70B [26] and GPT-3.5 on most benchmarks while being more computational efficient during both training and inference. Inspired by its strong performance, we follow its architectural design in the MoE layers and pre-train a MoE model in the protein domain.

## 3 Pre-training AIDO.Protein

To scale up model size while maintaining a high training and inference efficiency, we opt for sparse MoE architecture and pre-train a powerful protein language model with 16 billion parameters on a carefully curated protein sequence database.

### 3.1 Model architecture

As shown in Figure 1, our model is a transformer encoder-only architecture with the dense MLP layer in each transformer block replaced by a sparse MoE layer. The MoE layer design largely follows Mixtral 8×7B [19]. The MoE layer is applied independently for each token in the input sequence. To be specific, suppose we have *N* experts {*E*_*i*_(*x*), *i* = 1, …*N* − 1}, the output of the MoE layer *y* for each input token *x* is the weighted sum of the outputs of the expert networks, as shown in the following equation:

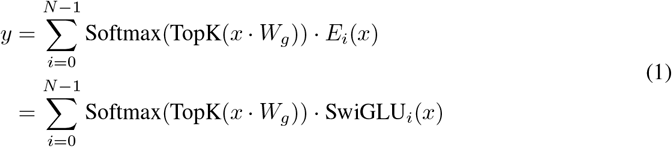

where *W*_*g*_ ∈ ℛ^(*d,N*)^ denotes the weight of the routing network (*d* is the hidden size), *E*_*i*_(*x*) = SwiGLU_*i*_(*x*) denotes the *i*-th expert netwok. In our experiment, we set *N* = 8, *K* = 2, *d* = 2304. Our model contains 36 transformer layers and 36 attention heads, totaling 16 billion parameters. During training and inference, each input token is processed by 4.5 billion parameters, using only 28% of the total number of parameters.

**Figure 1:**
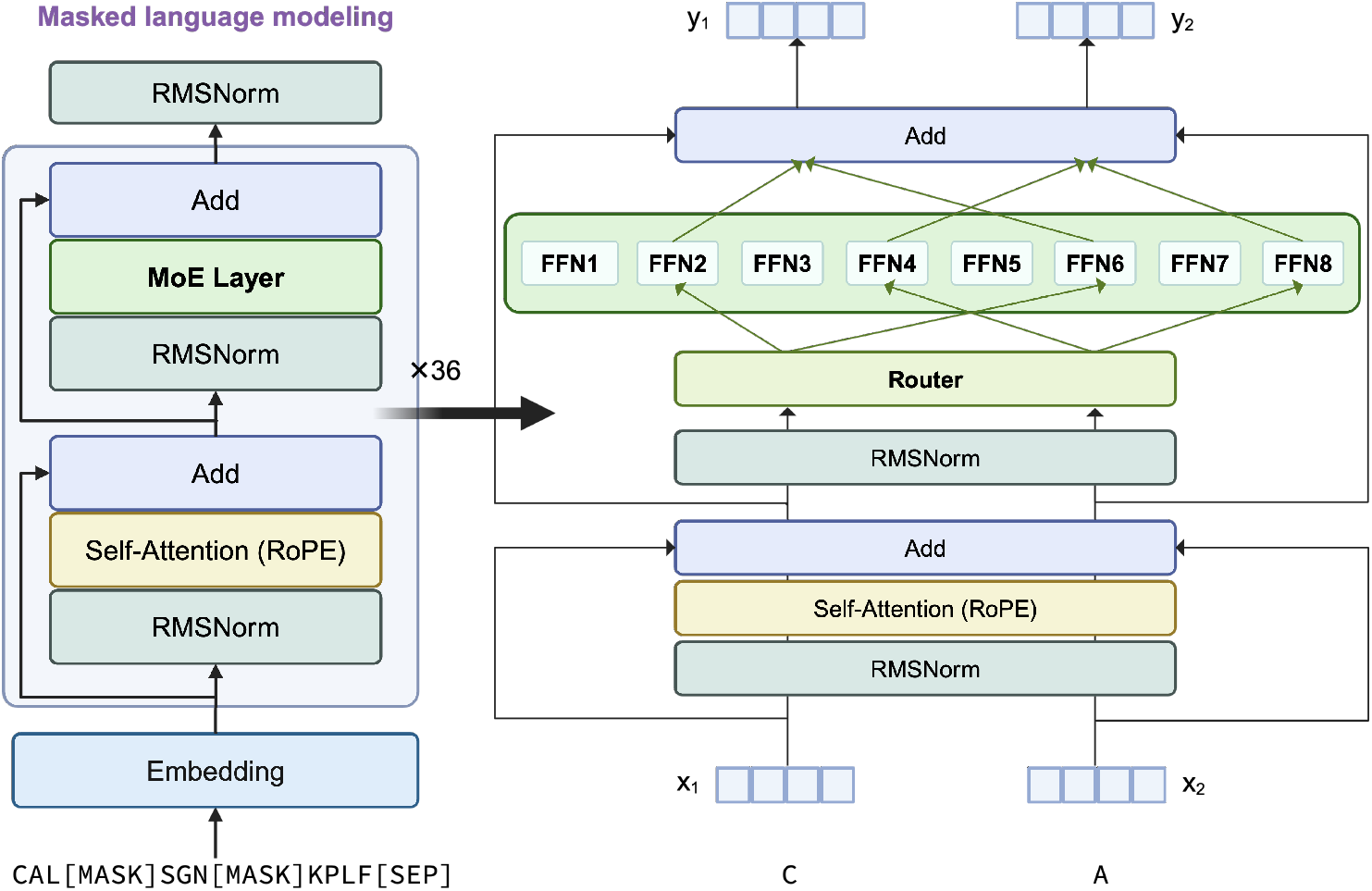
Model architecture of AIDO.Protein. We use sparse MoE architecture, with 8 experts in the Feed-Forward layer of a transformer block. For each token, 2 experts will be selectively activated by the top-2 routing mechanism. Figure created in BioRender.com.

AIDO.Protein is trained using the standard masked language modeling (MLM) objective. During training, the model predicts masked amino acids in a sequence, allowing it to learn the complex dependencies and relationships inherent in protein sequences. The use of MoE layers allows the model to allocate different experts to different types of sequence patterns, thus capturing a broader range of sequence features and enhancing its ability to predict and understand protein functions.

### 3.2 Pre-training data

Inspired by previous work [27], We initially pre-trained our model on the combination of Colab- foldDB [28] and UniRef90 [29] databases. The ColabfoldDB emphasizes metagenomic data sources such as BFD [30], MGnify [31], and various eukaryotic and viral datasets, including SMAG [32], MetaEuk [33], TOPAZ [34], MGV [35], and GPD [36]. UniRef90 offers clustered sets of sequences from the UniProt Knowledgebase ^3^ and selected UniParc ^4^ records to achieve comprehensive coverage of the sequence space at multiple resolutions while minimizing data redundancy. Specifically, we utilize UniRef90/50 (before December 2022), which includes incremental data beyond the UniRef50/S representatives.

Given the effectiveness of UniRef90 for previous protein language models [37, 38, 3], and the observed benefits of continuous training on domain-specific data for enhancing downstream task performance[39], we further train on UniRef90 with an additional 100 billion tokens.

In summary, we developed two versions of AIDO.Protein: AIDO.Protein-16B trained on 1.2 trillion amino acids from ColabfoldDB and Uniref90, and AIDO.Protein-16B-v1 continuously trained on an additional 100 billion amino acids from Uniref90.

### 3.3 Pre-training settings

We use a global batch size of 2048 and context length of 2048. For optimizer, we use AdamW with weight decay of 0.1. The cosine learning rate schedule is employed with warmup ratio set to 2.5% of the total training steps. To accelerate training, we use FP16 mix precision training. We adopt Megatron-Deepspeed framework and pre-train our model using 256 Nvidia A100-80G GPUs for 25 days.

The pre-training process consists of three stages. In the first stage, the model was trained on 1 trillion tokens sampled from the UniRef90 and Colab databases over 18.5 days, with a learning rate starting at 2e-4 and decaying to 2e-6. The second stage involved training on 200 billion tokens from the same data sources for 4 days, with the learning rate decreasing from 1e-5 to 1e-6. In the final stage, the model was further trained on 100 billion tokens from UniRef90, with a learning rate ranging between 1e-5 and 1e-6 for 2.5 days.

## 4 Experiments

We evaluate AIDO.Protein across more than 300 tasks from two important protein benchmarks, i.e., xTrimoPGLM benchmark [4] and ProteinGym DMS benchmark [40], encompassing residue-level, sequence-level, and protein-protein interaction (PPI) level tasks. Additionally, we adopt AIDO.Protein to develop a diffusion modeling framework for protein sequence design.

### 4.1 AIDO.Protein achieves strong results across diverse tasks from xTrimoPGLM benchmark

#### Tasks

To fully test our model’s ability in various protein understanding tasks, we evaluate our model on xTrimoPGLM benchmark [4]. It contains 18 diverse tasks which can be classified into four categories as follows:

- Protein structure prediction: **(1) Contact map prediction** aims to predict whether two residues *i* and *j* in a protein sequence are in contact or not based on their distance in the 3D structure with a threshold of 8Å. **(2) Fold prediction** aims to classify the protein sequence into one of the 1,195 known folds. **(3) Secondary structure prediction** aims to classify each residue into one of the 3 secondary structures, including Helix, Strand, and Coil.
- Protein function prediction: **(4) Antibiotic resistance prediction** aims to classify a protein sequence into one of the 19 antibiotics it is resistant to. **(5) Fluorescence prediction** aims to predict the fluorescence intensity of green fluorescent protein mutants. **(6) Fitness pre-diction** aims to to predict the fitness of GB1 binding following mutations. **(7) Localization prediction** aims to classify a protein sequence into one of the 10 subcellular localization categories.
- Protein-protein interaction prediction: **(8) Enzyme catalytic efficiency** aims to predict the enzymatic turnover numbers denoting the maximum chemical conversion rate of a reaction for a metabolic enzyme. **(9) Peptide-HLA/MHC affinity** aims to predict whether a given paired peptide and human leukocyte antigen (HLA) sequence can bind or not. **(10) Metal ion binding** aims to predict the existence of metal-ion binding site(s) on a given protein sequence. **(11) TCR pMHC affinity** aims to predict whether a given paired T cell receptor (TCR) sequence and peptide can bind or not.
- Protein development: **(12) Solubility** aims to predict whether a protein is soluble or insoluble. **(13) Stability** aims to predict the concentration of protease at which a protein can retain its folded state. **(14) Temperature stability** aims to predict a protein’s capacity to preserve its structural stability under temperature 65 degree Celsius. **(15) Optimal temperature** aims to predict the optimal temperature for the catalytic activity of an enzyme. **(16) Optimal ph** aims to predict the optimal pH for the enzyme’s reactions. **(17) Cloning clf** aims to predict whether a protein sequence tends to be a cloning failure or not. **(18) Material production** aims to predict whether a protein sequence fails at the protein material stage or not.

#### Fine-tuning models

We use LoRA [41] for efficient finetuning on the 18 tasks. For sequence-level classification/regression tasks, for each input protein sequence ^5^, we perform mean pooling over the output hidden states of the transformer encoder and use a two-layer MLP network as the prediction head. For the contact map prediction, a token-level pairwise classification task, we first compute the outer product for the output hidden states of the transformer encoder to obtain a feature map, and then use a 2-layer MLP with inter hidden size 128 for prediction. For the secondary structure prediction, a token-level classification task, we use a two-layer MLP as the prediction head with the inter hidden size set to 128.

#### Fine-tuning settings

We follow the train/valid/test splits and evaluation metrics in the xTrimoPGLM benchmark ^6^. For those tasks without validation sets, we randomly split 10% of training data for validation. For all tasks, we use LoRA fine-tuning with rank 16 and alpha 16. We use Adam optimizer with a peak learning rate 1e-4 and cosine learning rate scheduler with a warmup ratio of 0.05 for most of the tasks. For contact map prediction, we use Adam with a constant learning rate of 1e-4. We fine-tune the model for 10, 15 or 20 epochs and select the best checkpoints based on the validation scores. For details of hyperparameters for each tasks, please refer to our codebase.

#### Results

Table 1 shows the results of our model on the xTrimoPGLM benchmark. On 14 out of 18 tasks, our model AIDO.Protein-16B achieves better results than ESM2-15B, demonstrating that our sparse MoE model is effective while being more efficient in both training and testing. On average across the 18 tasks, we achieve a average score of 0.846, outperforming both ESM2-15B and xTrimoPGLM-100B. In particular, our model excels in protein structure prediction and protein development tasks, indicating that it can serve a powerful foundation model for protein design.

**Table 1:** AIDO.Protein-16B outperforms ESM2-15B on 14 out of 18 diverse tasks from the xTrimoPGLM benchmark. The results of ESM2-15B and xTrimoPGLM-100B are the LoRA finetuning results reported in [4]. Bold denotes the performance of our model is better than ESM2-15B.

| Cate. | Task | Metric | AIDO.Protein-16B (ours) | ESM2-15B | xTrimoPGLM-100B |
| --- | --- | --- | --- | --- | --- |
| Protein Structure | contact prediction | Top L/5 ACC | **0.925** | 0.922 | 0.933 |
|  | fold prediction | ACC | **0.763** | 0.692 | 0.756 |
|  | secondary structure prediction | ACC | **0.874** | 0.759 | 0.753 |
| Protein Function | antibiotic resistance | ACC | 0.979 | 0.983 | 0.984 |
|  | fluorescence prediction | Spearman CC | **0.679** | 0.637 | 0.660 |
|  | fitness prediction | Spearman CC | **0.950** | 0.948 | 0.961 |
|  | localization prediction | ACC | 0.811 | 0.824 | 0.816 |
| Protein Interaction | enzyme catalytic efficiency | Pearson CC | **0.749** | 0.746 | 0.748 |
|  | metal ion binding | ACC | **0.824** | 0.809 | 0.828 |
|  | peptide_HLA_MHC affinity | AUC | 0.971 | 0.973 | 0.967 |
|  | tcr_pmhc affinity | AUC | **0.944** | 0.941 | 0.951 |
| Protein Development | solubility prediction | ACC | **0.806** | 0.746 | 0.795 |
|  | stability prediction | Spearman CC | **0.824** | 0.808 | 0.842 |
|  | temperature stability | MCC | 0.932 | 0.932 | 0.942 |
|  | optimal temperature | Pearson CC | **0.798** | 0.733 | 0.740 |
|  | optimal ph | Spearman CC | **0.640** | 0.625 | 0.650 |
|  | cloning clf | AUC | **0.864** | 0.766 | 0.848 |
|  | material production | AUC | **0.888** | 0.792 | 0.865 |
| Average |  |  | **0.846** | 0.813 | 0.835 |

### 4.2 AIDO.Protein demonstrates impressive performance on ProteinGym DMS benchmark

We further evaluate AIDO.Protein on ProteinGym DMS benchmark to fully test our model’s ability in protein fitness prediction. This benchmark consists of 66 indels assays and 217 substitutions assays, with each assay providing all possible mutations for a specific target protein, along with their corresponding fitness scores. We adopt Spearman rank correlation and MSE as the evaluation metric. In the ProtienGym benchmarks, many methods are specialized and restricted to either indel or substitution tasks. We will focus mostly in comparing to methods which are versatile and generally applicable to both types of tasks. we will primarily compare our results with ESM-1v [37], Tranception [42], and MSA Transformer [43], with the latter two methods leveraging rich MSA information as input.

#### 4.2.1 DMS indels supervised benchmark

##### Tasks

Indel mutations are insertions or deletions of residues in a protein sequence. The DMS indels benchmark consists of 66 assays In machine learning, they can be formulated as sequence-level regression tasks. As shown in Table 2, the sample size for each task varies from 47 to 225,998, with *Q*_3_ = 193. In particular, 54 tasks in the indels benchmark have sample sizes smaller or equal to 205. When evaluating under the 5-fold cross-validation setting, the small sample size makes an expressive model prone to overfitting. Besides, the Spearnman rank correlation computed in a small validation set is not reliable for model selection. Therefore, we design different fine-tuning models for different tasks based on their sample sizes.

**Table 2:** Data statistics of ProteinGym DMS benchmark.

|  | Indels (66 assays) |  | Substitutions (217 assays) |  |
| --- | --- | --- | --- | --- |
|  | target seq len | sample size | target seq len | sample size |
| mean | 134 | 4,352 | 397 | 11,363 |
| std | 182 | 27,910 | 502 | 51,668 |
| min | 37 | 47 | 37 | 63 |
| 25% | 52 | 121 | 69 | 1,332 |
| 50% | 62 | 154 | 245 | 2,339 |
| 75% | 72 | 193 | 536 | 6,769 |
| max | 770 | 225,998 | 3423 | 53,6962 |

##### Fine-tuning settings

Following the evaluation setting in ProteinGym, all the tasks are evaluated under a 5-fold cross-validation setting with the fold split in line with the *Random* cross-validation scheme provided by ProteinGym. For the 54 small tasks, we use linear probing with AIDO.Protein frozen to alleviate overfitting. The prediction head is a 2-layer MLP with the inter hidden size set to 128 and dropout rate set to 0.1. We use AdamW optimizer with a peak learning rate of 1e-3 and cosine learning scheduler with warmup ratio of 0.05. We do not use a validation set for model selection. Instead, we directly train the model to 1,000 steps and then use the last checkpoint to predict the test labels. For the other 12 tasks which contain more samples, we use LoRA fine-tuning with rank 16, alpha 32, and peak learning rate 1e-4. We use one fold for validation and train the model to 10,000 steps with early stopping. For all the tasks, the batch size *B* is determined by the sample size *N* using following rules: if *N* ≤ 100, then *B* = 4 ; if 100 *< N* ≤ 1000, then *B* = 8; if 1000 *< N* ≤ 5000, then *B* = 16; if 5000 *< N* ≤ 10000, then *B* = 32; if *N >* 10000, then *B* = 64.

##### Results

As shown in the upper section of Table 3, our model achieves 0.748 corrected average Spearman correlation across 66 indels assays, ranking in the second place across all models in the leaderboard ^7^. Notably, the results of our model in terms of both mean square error (MSE) and Spearman correlation are very close to the SOTA model ESM-1v Embeddings [37]. And it achieves SOTA results in Fitness prediction, outperforming other models by large margins. Interestingly, our model outperforms MSA Transformer Embedding [43] and Tranception Embeddings [42], models that leverage homologous sequences for inference. This result indicates that protein language model is better at handling sequences with insertion and deletions while MSA models are not robust enough in this case. Detailed performance for each indel task is available in the Supplementary Figure 2.

**Table 3:** Results on ProteinGym DMS supervised benchmark. Bold denotes the best results, and underline denotes the second best results. AIDO.Protein achieves nearly 99% of the Spearman performance and superior MSE performance compared to Tranception Embeddings, the top MSA- based model in the overall DMS supervised benchmark, while significantly outperforming the previous state-of-the-art single-sequence model, ESM-1v.

| Task Type | Model | Spearman by Functions | | | | | Avg. Spearman ↑ | Avg. MSE ↓ |
| --- | --- | --- | --- | --- | --- | --- | --- | --- |
|  |  | Activity | Expression | Fitness | Stability | Binding |  |  |
| Indels<br>66 tasks | ESM-1v Embeddings | **0.706** | <u>0.729</u> | <u>0.718</u> | **0.856** | / | **0.752** | **0.365** |
|  | Tranception Embeddings | 0.674 | **0.753** | 0.716 | 0.797 | / | 0.735 | 0.410 |
|  | MSA Transformer Embeddings | 0.661 | 0.614 | 0.710 | 0.769 | / | 0.689 | 0.486 |
|  | AIDO.Protein (ours) | <u>0.698</u> | 0.721 | **0.742** | <u>0.829</u> | / | <u>0.748</u> | <u>0.370</u> |
| Substitutions<br>217 tasks | ESM-1v Embeddings | 0.559 | 0.641 | 0.534 | 0.634 | **0.881** | 0.639 | 0.563 |
|  | Tranception Embeddings | **0.615** | **0.716** | **0.610** | 0.872 | 0.672 | **0.696** | **0.503** |
|  | MSA Transformer Embeddings | <u>0.596</u> | 0.632 | 0.523 | <u>0.886</u> | 0.564 | 0.642 | 0.573 |
|  | AIDO.Protein (ours) | 0.574 | <u>0.677</u> | <u>0.569</u> | **0.913** | <u>0.675</u> | <u>0.682</u> | <u>0.509</u> |
| Overall<br>283 tasks | ESM-1v Embeddings | 0.593 | 0.662 | 0.577 | 0.686 | **0.881** | 0.665 | 0.517 |
|  | Tranception Embeddings | **0.629** | **0.725** | **0.635** | 0.855 | 0.672 | **0.705** | <u>0.481</u> |
|  | MSA Transformer Embeddings | <u>0.611</u> | 0.628 | 0.567 | <u>0.859</u> | 0.564 | 0.653 | 0.553 |
|  | AIDO.Protein (ours) | 0.603 | <u>0.687</u> | <u>0.609</u> | **0.893** | <u>0.675</u> | <u>0.697</u> | **0.477** |

**Figure 2:**
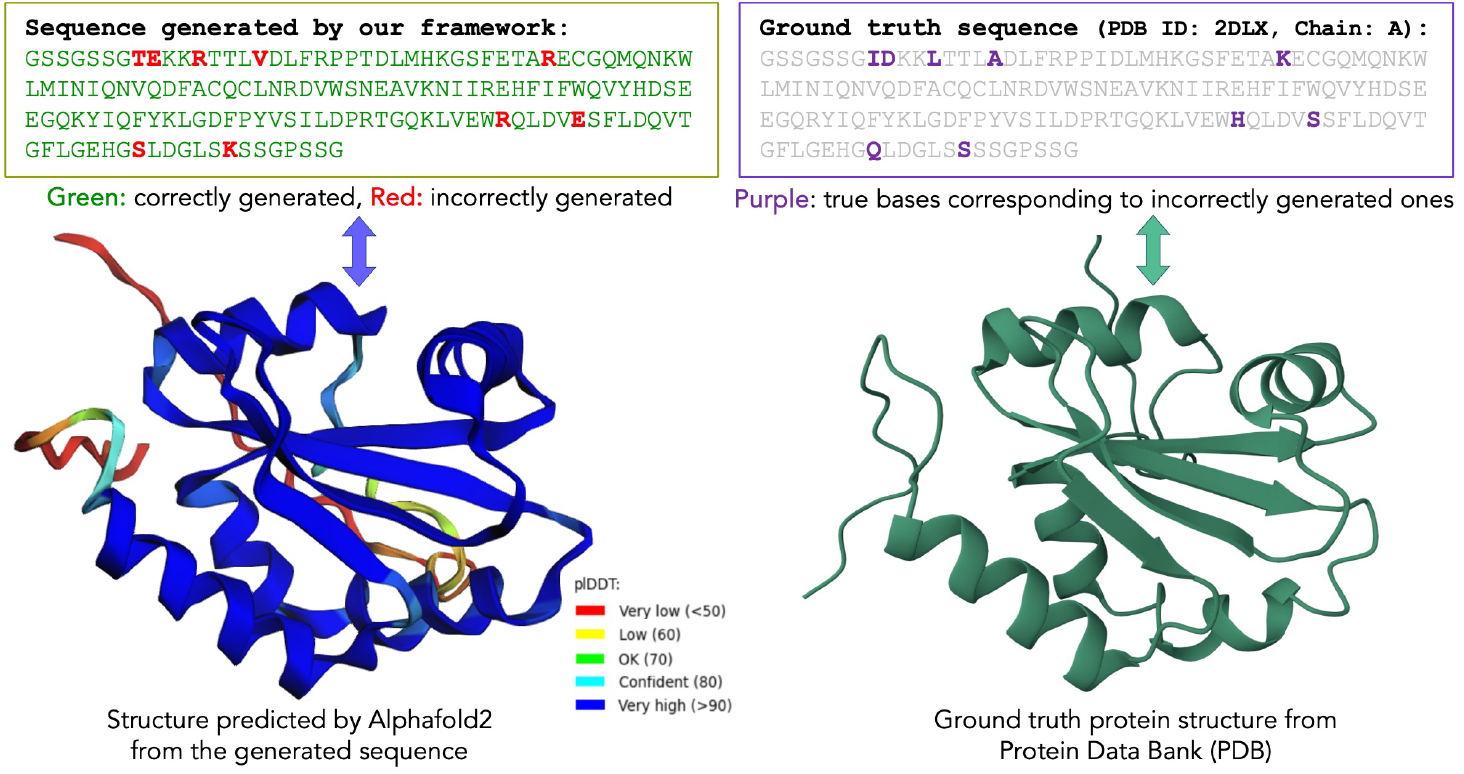
An example result of our protein inverse folding framework. (PDB ID: 2DLX, Chain: A). **Left:** Our discrete diffusion based inverse folding framework generates the protein sequence. We then use AlphaFold2 [47] to predict its structure from the generated sequence for analysis. The confidence of structure prediction, measured with pIDDT [47], is shown with color-codes. We use ColabFold [48] framework for AlphaFold2-based inference and rendering the protein molecule. **Right:** Ground truth protein sequence and structure from Protein Data Bank [49].

#### 4.2.2 DMS substitutions supervised benchmark

##### Tasks

Substitution mutations involve replacing one or more residues in a protein sequence with different ones. The substitution benchmark includes 217 assays, comprising 69 single substitution assays and 148 multiple substitution assays. As shown in Table 2, the sequence length ranges from 37 to 3423, with sample size varying from 63 to 536962. For tasks with small sample sizes, the model is prone to overfitting. Additionally, for tasks with excessively long sequences, fine-tuning the model leads to OOM issues. Therefore, we apply different finetuning strategies for different tasks based on the sample size and sequence length.

##### Finetuning settings

The ablation study in Supplementary Tab 1 shows that continuous pretraining on Uniref90 has significantly improved zero-shot substitution prediction, demonstrating the advantage of continuous training for substitution prediction. Therefore, we focus on evaluating AIDO.Protein-16B-v1, the version of AIDO.Protein-16B continuously trained on 100 billion amino acids from Uniref90, on the supervised substitution benchmark. ProteinGym provides three cross-validation schemes for the substitutions benchmark. To complete all tasks within reasonable time and resource constraints, we opted for the *Random* five-fold cross-validation scheme, consistent with the scheme used in the indels benchmark. We utilize the Adam optimizer and a cosine learning rate schedule. We employ LoRA [41] fine-tuning with a rank of 16 and alpha of 32, setting the peak learning rate to 1e-4 and training for 10,000 steps. Early stopping is triggered when the Spearman score on the validation set does not improve for predefined patience threshold. For 13 tasks with sample sizes exceeding 20,000, only one epoch per fold is performed, as a single pass through the data is sufficient. For 4 tasks with sequence lengths over 2048, we truncate the sequences to 2048 and adjust LoRA rank to 4 and alpha to 8. The batch size *B* and the early stopping patience *P* are determined by the sample size *N* using following rules: if *N* ≤ 100, then *B* = 4, *P* = 10 ; if 100 *< N* ≤ 1000, then *B* = 8, *P* = 5; if 1000 *< N* ≤ 5000, then *B* = 16, *P* = 3; if 5000 *< N* ≤ 10000, then *B* = 32, *P* = 3; if *N >* 10000, then *B* = 64, *P* = 1.

##### Results

As shown in middle section of Tab 3, our model achieves an average Spearman correlation of 0.682 and an mean square error (MSE) of 0.509 in supervised substitution benchmark, significantly outperforming previously reported best single-sequence based method, ESM-1v Embeddings, at both the functional group level and in average scores. Despite not utilizing MSA information, our model outperforms most MSA-based methods, such as the MSA Transformer, showcasing its strong ability to capture protein sequence information at the residue level. Furthermore, when compared to Tranception Embeddings, the leading MSA-based model in the overall DMS supervised benchmark, our model achieves comparable performance in both Spearman correlation and MSE metrics, and even surpasses Tranception Embeddings in the Stability and Binding functional groups. Further details on performance for each substitution task are provided in the Appendix Figure 3. Based on these comparisons, a promising direction for our future work would be to incorporate MSA into AIDO.Protein model for further improvements.

**Figure 3:**
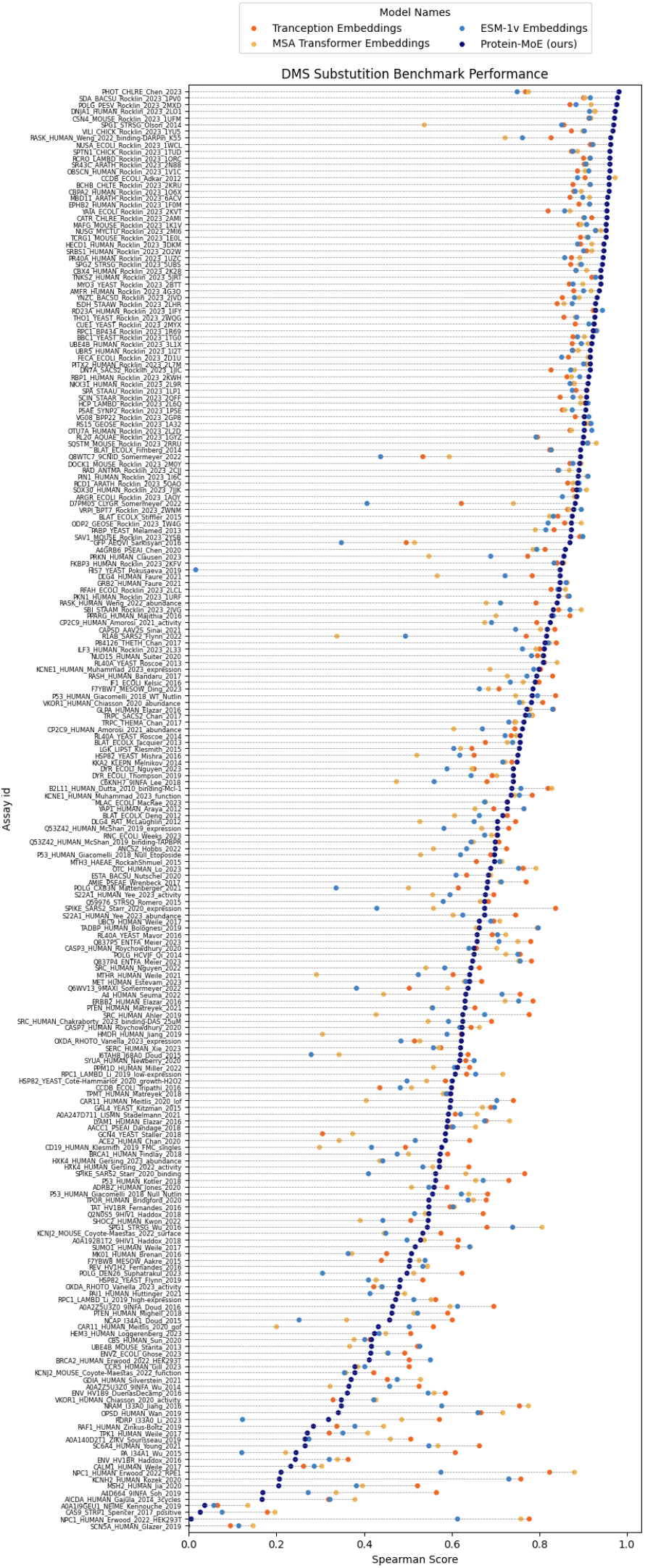
Spearman correlation performance on each assay from the substitution benchmark. We compared our AIDO.Protein (in dark blue) with the top 3 models in the overall DMS benchmark: Tranception (in orange) and MSA Transformer (in yellow), both of which are MSA-based, and ESM-1v (in light blue), which uses single sequence inputs. AIDO.Protein outperforms MSA Transformer and ESM-1v on most tasks, achieving performance close to Tranception.

**Figure 4:**
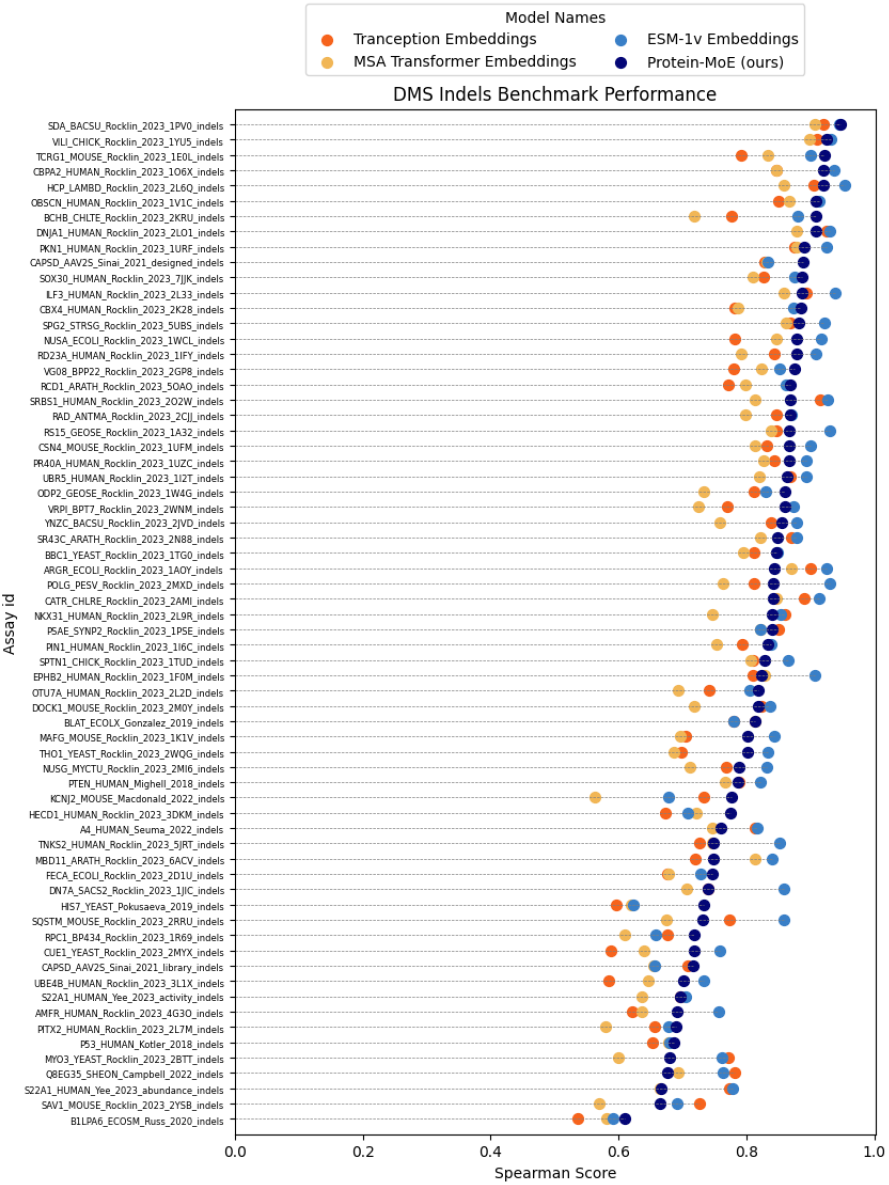
Spearman correlation performance on 66 indels assays from DMS indels benchmark. We compared our AIDO.Protein (in dark blue) with the top 3 models in the overall DMS benchmark: Tranception (in orange) and MSA Transformer (in yellow), both of which are MSA-based, and ESM-1v (in light blue), which uses single sequence inputs. AIDO.Protein outperforms both MSA based models on most tasks, achieving performance close to ESM-1v.

#### 4.2.3 DMS overall supervised benchmark

##### Results

In the bottom section of Tab 3, we finally evaluate our model’s overall performance across 66 indels and 217 substitutions tasks. Even without leveraging MSA, our model achieves nearly 99% of the average Spearman correlation and superior MSE performance compared to the overall best MSA-based model, Tranception Embeddings, and significantly outperforms the previously reported state-of-the-art model, ESM-1v Embeddings, which also does not use MSA.

### 4.3 AIDO.Protein offers enhanced capabilities for protein design

Protein design is a vital area of research and application in biochemistry, molecular biology, and biotechnology [50, 12]. This section discusses the adaptation of AIDO.Protein for this purpose. Specifically, we develop a discrete diffusion modeling [51, 52] framework for protein inverse folding, a crucial step in *de novo* protein design, as discussed by Mu *et al*.[53]. We adopt ProteinMPNN-CMLM, a variant of ProteinMPNN [20] produced by [21], as our structure encoder for structure-conditioned diffusion. Details about the framework are available in Appendix A.1.

For experiments, we use the dataset by CATH 4.2 dataset [54], a well-established resource for evaluating protein design. In Table 4, we show our framework, denoted as AIDO.ProteinIF, achieves significantly higher score than the previous state-of-the-art methods.

**Table 4:** Comparison of protein inverse folding performance. Our protein design framework, with AIDO.Protein as the backbone, surpasses the performances of existing fixed-backbone protein inverse folding methods. The recovery rates of previous methods are quoted from [44].

| Model | Median Sequence Recovery |
| --- | --- |
| StructTrans [45] | 35.82 % |
| GVP [46] | 39.47 % |
| ProteinMPNN [20] | 45.96 % |
| ProteinMPNN-CMLM [21] | 48.62 % |
| LM-Design [21] | 54.41 % |
| DPLM [44] | 54.54 % |
| AIDO.Protein-IFdiff (ours) | **58.26 %** |

In Figure 2, we show an example generated protein by our framework and the ground truth from PDB [49].

## 5 Conclusions and future work

In this paper, we introduce AIDO.Protein, a 16 billion parameter protein language model that incorporates mixture-of-expert layers and is trained on 1.2 trillion amino acids from ColabfoldDB and UniRef90. To our knowledge, our work is the first application of sparse expert models in the protein domain, allowing for efficient modeling while maintaining high performance. AIDO.Protein demonstrates exceptional performance across various protein understanding tasks, achieving state-of-the-art (SOTA) results on most xTrimoPGLM tasks and ranking second on the ProteinGym DMS leaderboard. Additionally, it showcases remarkable potential in de novo protein design through inverse folding. This dual proficiency in understanding and generating proteins underscores the model’s value in advancing drug discovery, personalized medicine, enzyme engineering, and immune response prediction. Its capabilities position AIDO.Protein as a catalyst for significant innovations in biotechnology and synthetic biology, paving the way for new solutions and applications in these critical fields.

## A Experiments

### A.1 Protein Design

#### A.1.1 Masked Diffusion for Protein Sequence Generation

We aim to approximate a data distribution *q*(*x*) by training a diffusion model, by first iteratively adding noise to a sample *x*_0_ ∼ *q*(*x*) for *T* discrete steps (forward process) that results in a sample with entire noise *x*_*T*_; and then training a model, parameterized by *θ*, to denoise *x*_*T*_ iteratively to retrieve the original signal *x*_0_ (reverse process). In case of continuous signal, such as image or audio, at any time-step *t* ∈ [0, *T*], the sample *x*_*t*_ can be assumed as a linear combination of the original signal *x*_0_ and Gaussian noise *ϵ* ∼ 𝒩(0, 1) [55],

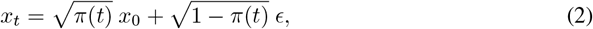

where *π*(.) ∈ [0, 1] is a monotonically decreasing function of time-step *t*. The model learns a marginal distribution *p*_*θ*_(*x*_*t−*1_|*x*_*t*_), which aims to approximate the true transition probability *q*(*x*_*t−*1_|*x*_*t*_, *x*_0_) of estimating a less noisy variant *x*_*t−*1_ given a relatively more noisy variant *x*_*t*_.

Given that, at *t* = 0 we have *x*_*t*_ = *x*_0_ (with *π*(*t*) = 1), and at *t* = *T, x*_*t*_ = *x*_*T*_ = *ϵ* ∼ 𝒩(0, 1) (with *π*(*t*) = 0) that is pure Gaussian noise. However, in case of discrete signals like protein sequence, this is infeasible to represent *x*_*T*_ as a samples from unit-Gaussian. We can, instead, represent *x*_*T*_ entirely by *absorbing state* [51, 56] that contain no data-specific signal, i.e., analogous to pure Gaussian noise. Following [56], we use [MASK] token as the absorbing state.

For our masked diffusion model training objective, we adopt the formulation proposed by [51]. Overall the objective function for diffusion, negative evidence lower bound on log likelihood (NELBO) [56], can be decoupled into three disjoint objectives for reconstruction 𝒥_*recon*_, diffusion 𝒥_*diff*_, and prior 𝒥_*prior*_. As derived by [56, 51, 52], it is possible to show that for diffusion directly on data samples *x*, 𝒥_*recon*_, 𝒥_*prior*_ = 0. Given this, NELBO for discrete times-steps *T* simplifies to,

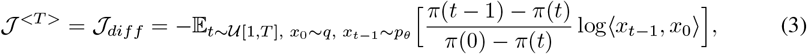

where 𝒰[1, *T*] is a uniform distribution integers between 1 and *T*, and ⟨*x*_*t*_−_1_, *x*_0_⟩computes the similarity between *x*_*t*−1_ and *x*_0_. Equation 3 is derived from the Kullback–Leibler divergence [57] between the transition probability distributions *q* and *p*. Please find the detailed derivation in [51].

In this study, we adopt cross-entropy loss between *x*_0_ and *x*_*t−*1_, ℒ_*CE*_(*x*_0_, *x*_*t−*1_), for − log⟨*x*_*t−*1_, *x*_0_⟩. [58] showed that, with higher number of diffusion steps T, we can get a tighter bound on 𝒥^*<T>*^. With *T* → ∞, Equation 3 becomes,

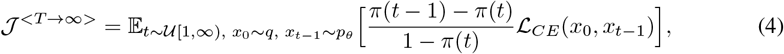

where *π*(0) = 1. Please note that, for *T* → ∞, *π*(*t* − 1) → *π*(*t*), i.e., the change in *π*(.) at any time *t* should be infinitesimally small. Also, since *π*(.) is monotonically decreasing, *π*(*t* − 1) − *π*(*t*) *>* 0. With *T* → ∞, we can represent this change with the negative time-derivative of *π*(.) at time *t*, 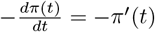. This leads to the continuous-time likelihood bound,

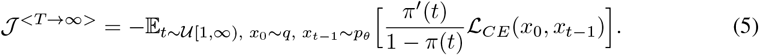

As shown by [51], the choice of *π*(.) has insignificant effect on the overall performance of the training algorithm. We adopt 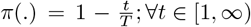 as our noise schedule. This further simplifies Equation 5 as,

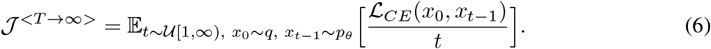

##### Intuition behind the objective function

Note that the loss computed on any sample *x*_*t*_ is now inversely proportional to *t*. Intuitively, if *t* is large, *x*_*t*_ is more noisy and hence it can potentially lead to many varieties of reconstructed samples 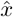 from *q*(*x*), i.e., all of them can be valid. However, with ℒ_*CE*_(*x*_0_, *x*_*t*_) loss we are always pushing the *x*_*t*_ to be more similar to *x*_0_, i.e., encouraging less diversity in generation, which is only expected if *x*_*t*_ is already very similar to *x*_0_ (when *t* is smaller). To address this conflict, the loss ℒ_*CE*_(*x*_0_, *x*_*t*_) is down-weighted by the factor 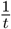.

#### A.1.2 Protein Inverse Folding

Protein inverse folding represents a cutting-edge computational technique aimed at generating protein sequences that will fold into specific three-dimensional structures [21, 20]. This innovative approach stands in stark contrast to traditional methods of protein folding, where the primary goal is to predict the 3D structure based on an existing protein sequence [2].

The central challenge in protein inverse folding involves identifying sequences capable of reliably adopting the intended structure [21, 44]. In our research, we concentrate on designing sequences based on the known backbone structure of a protein [21, 20, 44]. This is particularly crucial for fields like synthetic biology and nanotechnology, where the development of specific protein structures is essential for executing vital biological functions [50]. Recent advances in computational modeling, particularly those leveraging deep generative models, have significantly improved the accessibility and effectiveness of protein inverse folding approaches [21, 44, 20].

In the following part, we discuss how we adapt our AIDO.Protein with masked diffusion modeling conditioned on 3D protein structures.

##### Adaptation with Conditional Diffusion

During training, we aim to optimize the diffusion objective described in Equation 6. We start by sampling a sequence *x*_0_ with a known 3D protein structure and masking a fraction 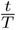 (where *t* ∼ 𝒰[1, *T*]). This results in *x*_*t*_, which acts as a noisy variant of *x*_0_.

We then pass *x*_*t*_ through ProteinMPNN-CMLM [21], generating an initial estimate of the sequence *S*_*t*_ and a structural embedding 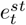.

This *S*_*t*_ is subsequently processed by the encoder of AIDO.Protein, producing the sequence embedding 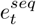. Following this, 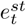 and 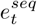 are input into an adaptor module [44, 21], which, in our design, consists of a multihead self-attention layer [59] combined with a bottleneck multi-layer perceptron [60], generating a new estimate of the protein sequence, *x*_*t−*1_. Note that the framework described above models the transition function *p*_*θ*_(*x*_*t−*1_|*x*_*t*_).

After the training is completed, we can generate sequences using our framework. We begin by providing the generation framework with a sequence composed solely of mask tokens, denoted by *x*_*T*_, along with the protein structure. The output *x*_*T*_−_1_ is anticipated to be a less noisy version of the expected ground truth *x*_0_. We then iteratively denoise this sequence over multiple steps to produce our final generated sequence 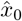.

### A.2 AIDO.Protein Performance on DMS Substitution and Indel Benchmarks

### A.3 Ablation Studies of Continuous Training

In this section, we further explore the impact of continuous training and how the choice of continuous training datasets affects downstream performance.

#### A.3.1 Influence of continuous training on model effectiveness

Previous studies [61, 39, 62] have shown that continuous training on domain-specific datasets can significantly improve performance on downstream tasks. We compare three models: the first is trained on 1 trillion tokens from UniMeta, the second continues with an additional 200 billion UniMeta tokens at a reduced learning rate, and the third focuses on UniRef90, a subset of UniMeta, with an additional 100 billion tokens. The performance of these three models on the DMS zero-shot benchmark is presented in Table 5. We observe that training on more UniMeta tokens significantly improves performance, and continuous training on a more domain-specific dataset, like UniRef90, further enhances results.

**Table 5:** The impact of continuous training on AIDO.Protein-16B performance in the DMS zero-shot benchmark.

| Model | # Total Tokens | Score |
| --- | --- | --- |
| AIDO.Protein-16B | 1 trillion | 0.401 |
| AIDO.Protein-16B | 1.2 trillion | 0.405 |
| AIDO.Protein-16B | 1.3 trillion | **0.407** |

#### A.3.2 Effects of continuous training dataset on model performance

We then discuss how the choice of continuous training datasets affects model performance. We continue training AIDO.Protein-16B, which has 1.2 trillion parameters, with an additional 100 billion tokens from three different datasets: UniRef90, UniRef50, and a sampled dataset combining UniRef90 and UniRef50. UniRef90 is a subset of UniMeta, while UniRef50 is a further clustered version of UniRef90. Inspired by ESM-2 [3], which proposes a sampling strategy to enhance data diversity and reduce redundancy by utilizing both UniRef90 and UniRef50, we adopt a similar approach to create the sampled dataset UniRef90/50. We randomly selected 50 zero-shot tasks from DMS zero-shot benchmark to compare the performance of these three continuous training models. The results are presented in Table 6. The model trained with Uniref90 achieves the best performance, while the model trained on both Uniref90 and Uniref50 outperforms the one solely trained on Uniref50, highlighting the benefits of utilizing Uniref90.

**Table 6:**
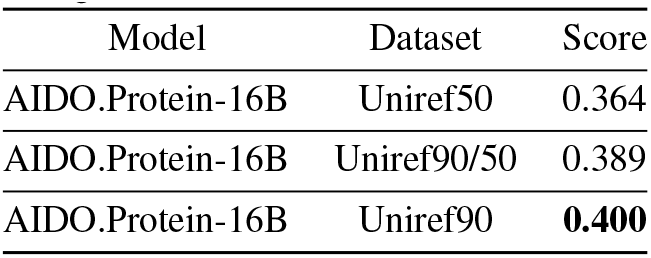
Comparison of AIDO.Protein with 1.3 trillion amino acids on selected DMS zeroshot tasks across various continuous training datasets.

| Model | Dataset | Score |
| --- | --- | --- |
| AIDO.Protein-16B | Uniref50 | 0.364 |
| AIDO.Protein-16B | Uniref90/50 | 0.389 |
| AIDO.Protein-16B | Uniref90 | **0.400** |

## B Data and Code Availability

We developed the ModelGenerator package to reproduce, apply, and extend the results in this manuscript https://github.com/genbio-ai/ModelGenerator.

Pre-trained models and data splits are also available on Hugging Face at https://huggingface.co/genbio-ai.

